# Analyzing the morphological variation of the Galápagos mistletoe *Phoradendron berteroanum* (Santalaceae) using herbarium specimens

**DOI:** 10.1101/2025.05.16.654492

**Authors:** Juliana Puentes-Marín, Anakah Denison, Justine Villalba-Alemán, Johny Mazón-Redín, C. Miguel Pinto

## Abstract

The hemiparasitic mistletoe *Phoradendron berteroanum* in the Galápagos shows remarkable morphological variation suggesting either the presence of clinal or abrupt geographical variation. To find patterns of the morphological variation of *P. berteroanum*. Eight vegetative and flower traits were measured from 68 herbarium specimens. Univariate and multivariate analyses comparing the traits by altitude, island, host and vegetation cover were performed. It was found that leaf size and internode length were larger at mid-elevations, while the number of floral segments decreased with altitude. Flower number varied by host, notably higher on *Scalesia baurii* from Pinzón Island. No evidence of unique island populations was found, so there is no morphological support for considering more than one species of mistletoe in the Galápagos. These results, however, reveal how *P. berteroanum* respond to local environmental conditions across the Galápagos.

## INTRODUCTION

The order Santalales is the largest lineage of parasitic flowering plants, comprising approximately 2,500 species with diverse growth habits, including herbs, trees, shrubs, and lianas (Nickrent et al., 2010; Su et al., 2015). Members of this order are hemiparasitic, either facultative or obligate, and can parasitize roots or branches. Parasitism on branches is referred to as aerial parasitism, and plants exhibiting this condition are commonly known as mistletoes. Globally, mistletoes are classified into five families, 87 genera, and about 1,700 species, with Loranthaceae and Viscaceae sensu lato (including *Phoradendron* under Santalaceae in the current classification) being the most diverse (Amico et al., 2019; Nickrent et al., 2010; Tinoco□Domínguez et al., 2024). All members of Viscaceae are aerial hemiparasites, and their fruits are dispersed by birds (Dettke and Caires, 2021; Kuijt, 2003; Kuijt & Hansen, 2015).

*Phoradendron* Nutt. with 278 described species is the largest genus of mistletoes in the Americas (Kuijt, 2003; Kuijt & Hansen, 2015). Among them, *Phoradendron berteroanum* (DC.) Griseb. has a wide range of distribution. It occurs in Central America (Costa Rica and Panama), the Caribbean (Cuba, Dominican Republic, Haiti, Jamaica and Puerto Rico), and several South American countries: Bolivia, Brazil, Colombia, Venezuela and Ecuador, including the Galápagos Islands. The numerous changes in the taxonomy of *P. berteroanum* over the years highlight its remarkable morphological variation, as observed in the Galápagos Islands (Kuijt, 2003; Wiggins et al., 1971). This morphological variability has often been interpreted as representing distinct species, leading to the application of different names, such as *P. henslovii* (Hook.f.) B.L.Rob., *P. galapageium* (Hook.f.) B.L.Rob., and *P. uncinatum* B.L.Rob. (Wiggins et al., 1971). However, these names are now considered synonyms of *P. berteroanum* (Kuijt, 2003; McMullen, 1999), reflecting on the adaptability—and phenotypic plasticity—of a single species to the diverse environments across the archipelago.

In the Galápagos Islands, *Phoradendron berteroanum* is found across a broad altitudinal range, from 25 to 1590 meters above sea level, inhabiting both the moist highlands and the arid lowlands, including volcanic rocky outcrops. We tested the hypothesis that its considerable morphological diversity in the archipelago reflects clinal variation—traits change gradually along environmental gradients. To do so, we examined eight morphological traits from herbarium specimens and analyzed their variation with respect to island, host species, and elevation.

## MATERIALS AND METHODS

### Data acquisition

A total of 68 herbarium specimens of *Phoradendron berteroanum*, collected between 1963 and 2022 and stored at the Charles Darwin Research Station (CDS) Herbarium, were analyzed (Figure 1). These specimens originate from eight islands of the Galápagos Archipelago: Santa Cruz, Isabela, Floreana, Pinta, Santiago, San Cristóbal, Pinzón, and Fernandina; these span an altitudinal range from 25 to 1,590 meters above sea level (masl). Land cover classes of the collecting sites were assigned via spatial intersection with a GIS layer developed by Rivas-Torres et al. (2018) (Figure 2). However, it is important to note that the number of specimens per island is uneven, with some islands being more represented than others. This discrepancy in sampling effort may introduce bias and should be considered when interpreting island-level patterns. The data supporting the findings of this study are available in the Charles Darwin Foundations repository: https://researchdata.fcdarwin.org.ec/records/202cw-xhy96

**Figure 1.**
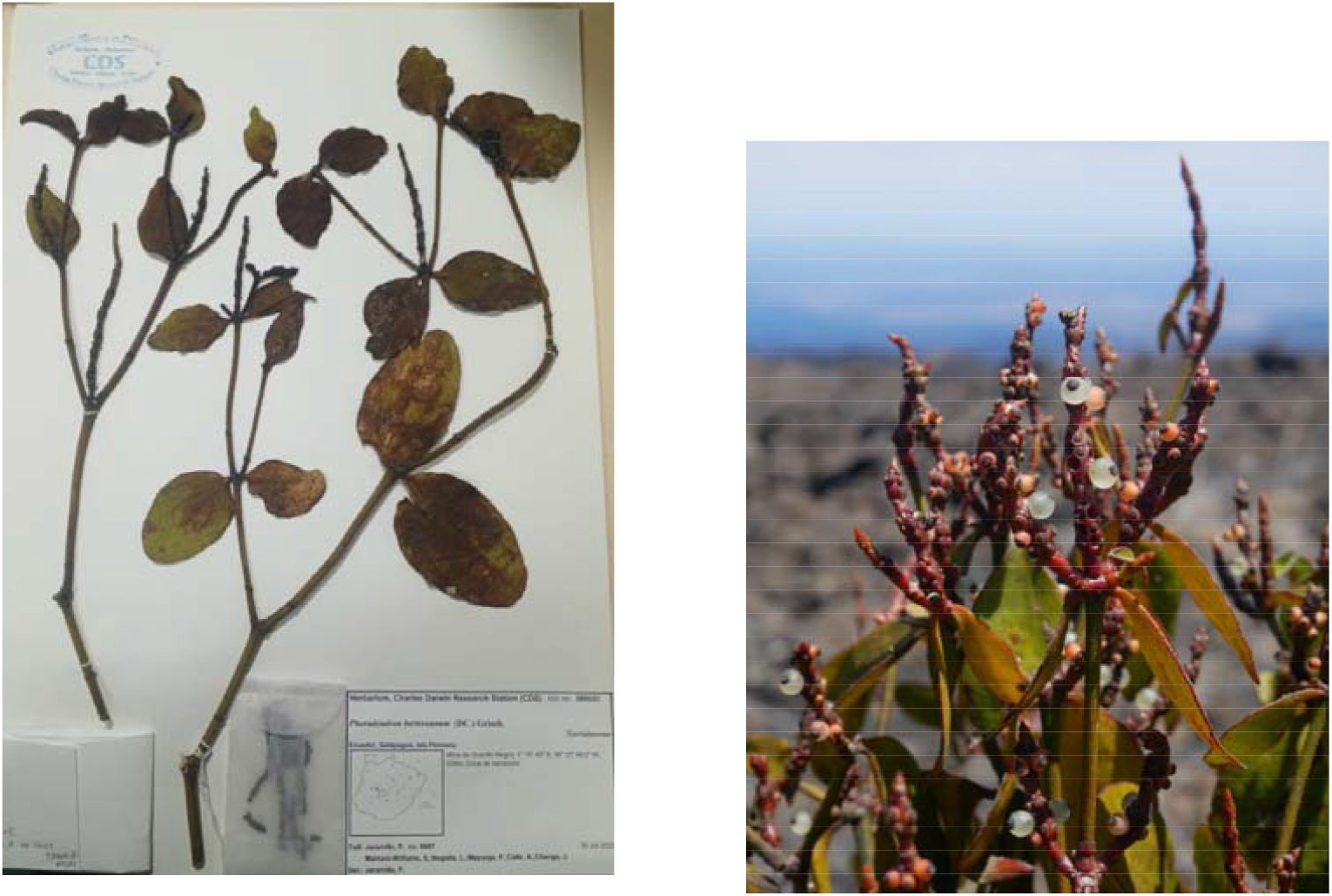
*Phoradendron berteroanum*. Left: Herbarium specimen CDS accession number: 58602; photo by Juliana Puentes-Marín. Right: *Phoradendron berteroanum* individual in the wild, in Sierra Negra Volcano, Isabela Island; photo by: Miguel Andrade.

**Figure 2.**
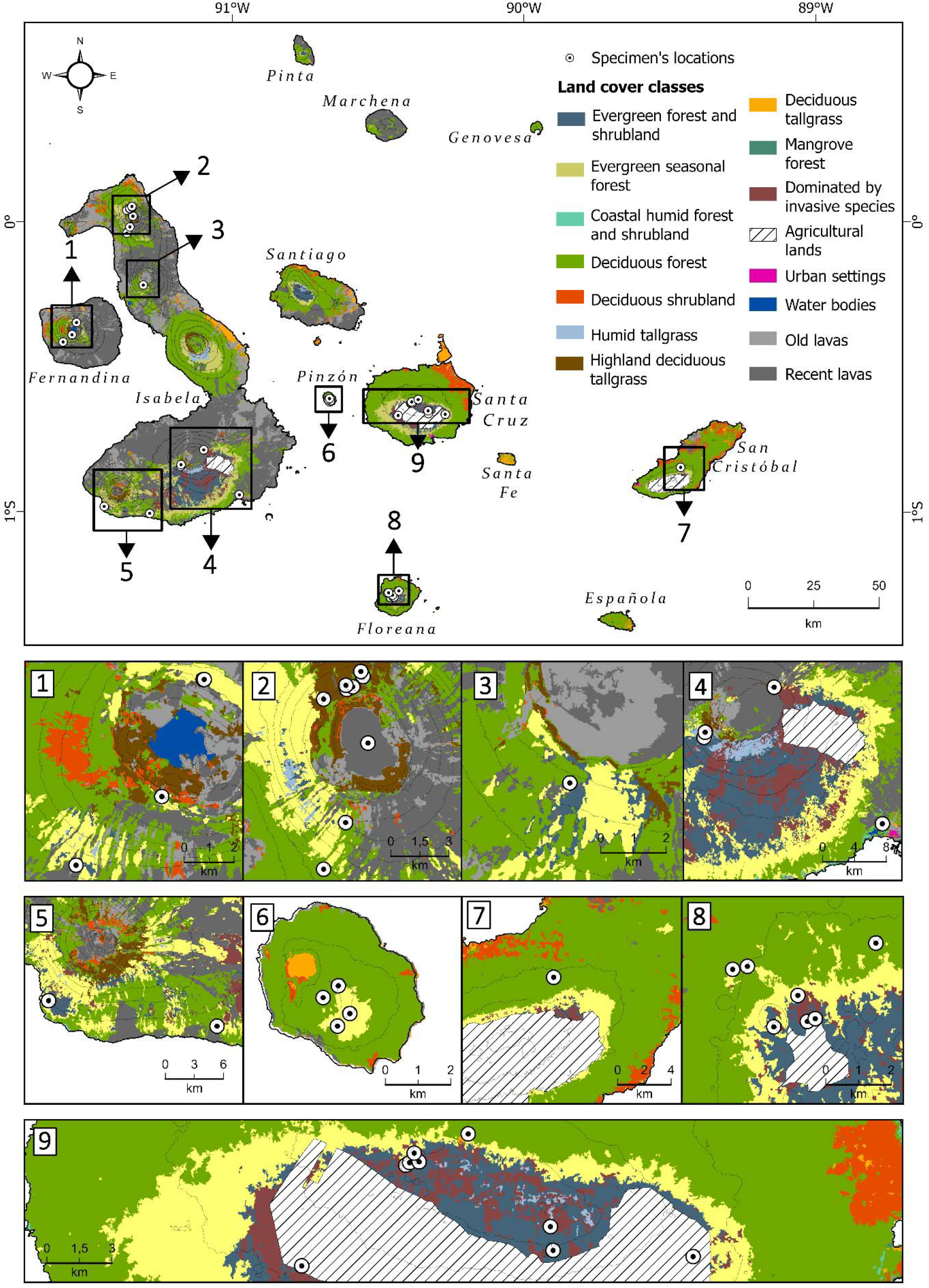
*Phoradendron berteroanum* distribution map by sampling localities. Specimens without coordinates were not included in the figure.

Eight morphological traits, focusing on leaf characteristics, internode morphology and inflorescence structure, were measured from each specimen. These morphological traits are: i) leaf length (LL), ii) leaf width (LW), iii) internode length (IN), iv) infertile segments number (NI), v) infertile segment length (IL), vi) floral segments number (FS), vii) floral segment length (FL), and viii) flower number (FN). For spikes that were glued down, the number of visible sockets on the accessible half was counted, and the total was estimated based on the observed pattern and the size of the spikes. Measurements were taken manually using a ruler, assisted by a table-mounted magnifying lens. Due to the preservative nature of herbaria, many standard measurements—such as specific leaf area (SLA)—could not be obtained. Most of the specimens were already mounted, lacked records of fresh weights, and were extremely delicate. LL was measured excluding the petiole, and LW was taken at the widest point. Segment length (IL and FL) were measured from the lowest visible point (inside the chevron joining point) to the analogous corner at the top edge, or to the tip in the case of terminal segments. IN was measured between nodes, from the innermost edge of the surface bract or scar. Additionally, the collection site, altitude, and host species recorded on the herbarium labels were documented for each specimen (Table S1). Outliers in the data were checked and corroborated ensuring that they weren’t procedural errors in measurement. The distribution of the data was analyzed using the fitdistr() function, from the package MASS. Lognormal transformations were applied to the FS and IN variables, since they did display a normal distribution, unlike the other parameters involved.

### Morphometric analyses

A principal component analysis (PCA) and a linear discriminant analysis (LDA) were performed. PCA was carried out using the prcomp() function from the stats package with a correlation matrix, while LDA was conducted using the lda() function from the MASS package. Specimens were grouped by elevation range (low, medium, and high) to explore potential patterns and assess whether distinct groupings emerged across altitudinal gradients. Subsequently, ANOVA tests were performed to compare the mean values of each morphological trait across categories of elevation, island, host species, and land cover type. Analyses were conducted using the aov() function, followed by Tukey’s post hoc comparisons via the TukeyHSD() function. It should be noted that when categorizing specimens by island or land cover type, the number of observations per group varies considerably, which may influence the comparability between categories and should be taken into account when interpreting the results. All statistical analyses were carried out in R software version 4.4.2(R Core Team, n.d.).

### Host specificity

To estimate the diversity of host species used by *Phoradendron berteroanum*, the host specificity index (KQ) was calculated (Amico et al., 2019; Kavanagh and Burns, 2012). The KQ index accounts for both the number of distinct host species and the frequency of collections from each host. It is calculated using the formula: 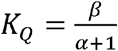, where **β** represents the minimum host range (i.e., the number of different host species recorded), and **α** is the sum of redundant collection records of all hosts. The KQ index is analogous to other diversity metrics, such as the Shannon index, as it incorporates both species richness and evenness in host usage (Kavanagh & Burns, 2012). This index was used to assess host specificity in *P. berteroanum* at different altitudinal bands (low, middle and high elevation). To assess whether morphological variability in *P. berteroanum* is associated with its host species, an ANOVA was conducted for each morphological trait, comparing trait values across the different identified host species.

## RESULTS

For the morphological traits in *Phoradendron berteroanum* herbarium specimens measured, the mean leaf length (LL) was 6.00 ± 1.85 cm, and the mean leaf width (LW) was 2.76 ± 0.94 cm. The mean internode length (IN) was 6.97 ± 2.41 cm. The mean number of infertile segments (NI) was 1.48 ± 0.49, with a mean infertile segment length (IL) of 0.29 ± 0.10 cm. Regarding reproductive traits, the mean number of floral segments (FS) was 4.83 ± 1.01, their mean length (FL) was 0.72 ± 0.42 cm, and the mean number of flowers (FN) per inflorescence was 16.50 ± 6.88.

### Morphological traits

The PCA and LDA analyses show no particular clustering by geographic or altitudinal distribution with respect to the morphometric traits measured (Figure 3). Likewise, there was no significant correlation between the traits of the specimens and their elevation, with all Pearson correlation coefficients lower than 0.01. However, when altitudinal elevation is categorized into 3 groups—low (0m – 499m), medium (500m – 999m) and high (above 1000m)—ANOVA results indicate significant differences in some of these traits among the different elevation groups (Figure 4).

**Figure 3.**
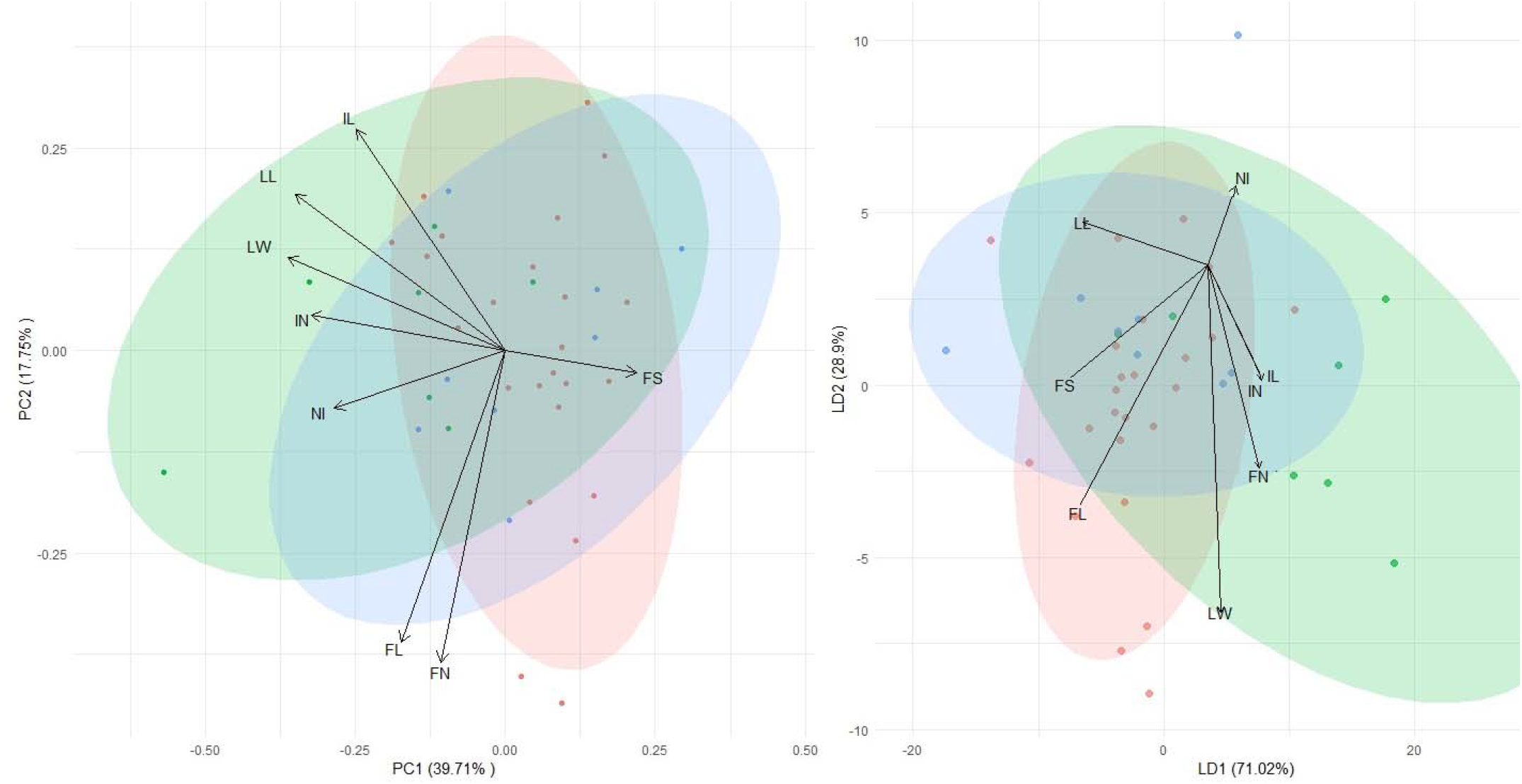
Left: principal component analysis. Right: linear discriminant analysis showing the associations between *P. berteroanum* specimens

**Figure 4.**
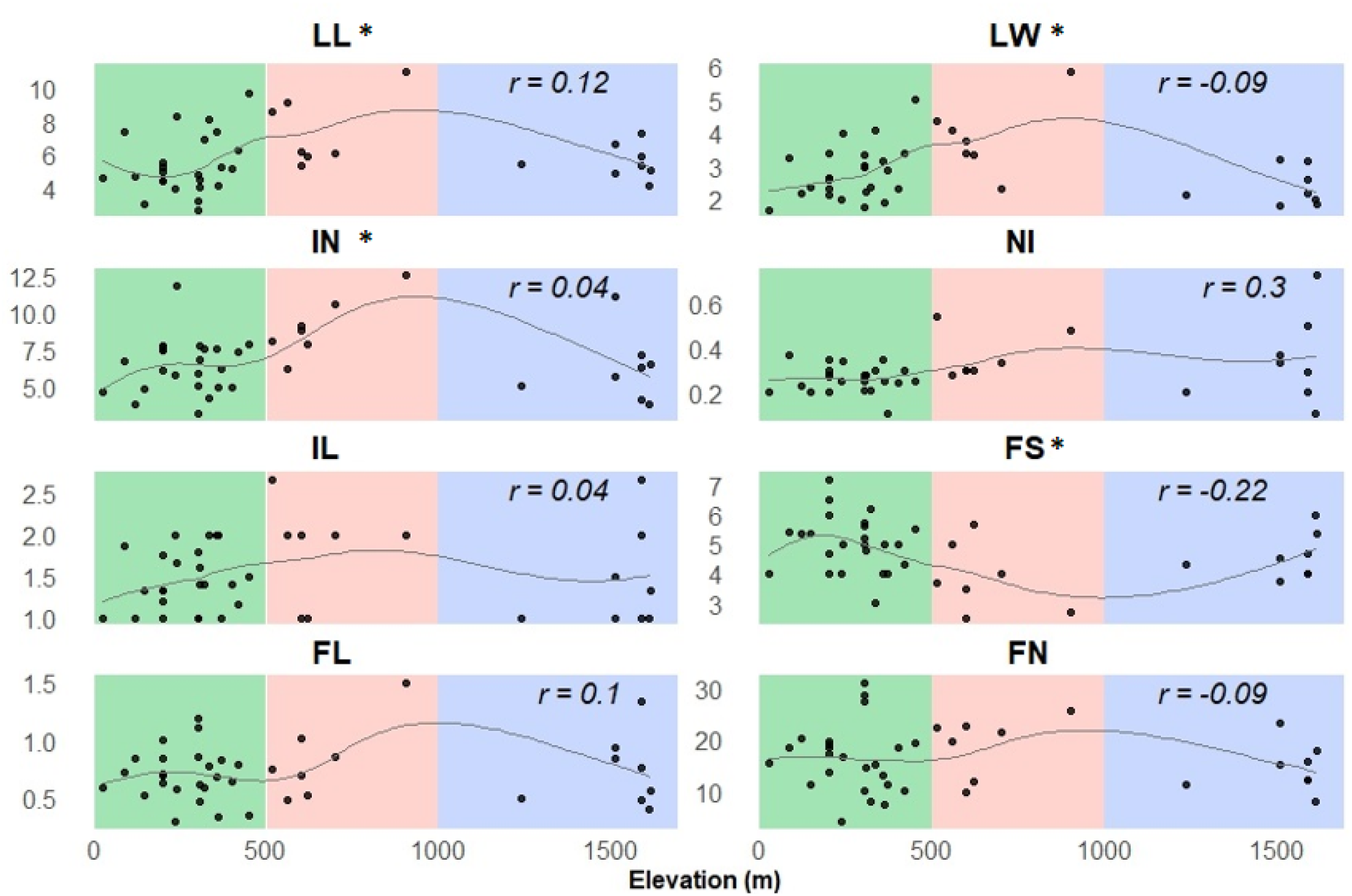
Scatterplots showing variation of *Phoradendron berteroanum* morphometric traits across three altitudinal ranges: Low (0-499 masl), Medium (500-999 masl) and High (above 1000 masl). Asterisks (*) indicate statistically significant trait differences between the altitude categories observed after conducting ANOVA test (p < 0.001); *r* values were obtained after conducting Pearsons’s correlation test. Trend curves were fitted using the LOESS (Locally Estimated Scatterplot Smoothing) method. Leaf length (LL), leaf width (LW), internode length (IN), infertile segments number (NI), infertile segment length (IL), floral segments number (FS), floral segment length (FL), flower number (FN).

Morphological traits related to the leaves and internodes of *Phoradendron berteroanum* show significant variation along the altitudinal gradient in the Galápagos Islands. Leaf length (LL, *p* = 0.0249) and leaf width (LW, *p* = 0.00316) reach their highest values at mid-elevations, whereas lower and higher elevations are associated with smaller measurements. Similarly, internode length (IN, *p* = 0.00917) is also greater at mid-elevation, with lower and more homogeneous values observed at the extremes. In contrast, the length of infertile segments (IL, *p* = 0.022) follows a distinct pattern, reaching its lowest values around mid-elevations. (Figure 4).

The ANOVA results indicate that leaf length (LL) and leaf width (LW) vary significantly among islands, whereas the remaining morphological traits show no significant differences. For LL, significant differences are found between Pinta and Isabela (*p* = 0.0273) and between Pinzón and Pinta (*p* = 0.0014). Similarly, for LW, a significant difference is observed between Pinta and Isabela (*p* = 0.0310) (Figure 5).

**Figure 5.**
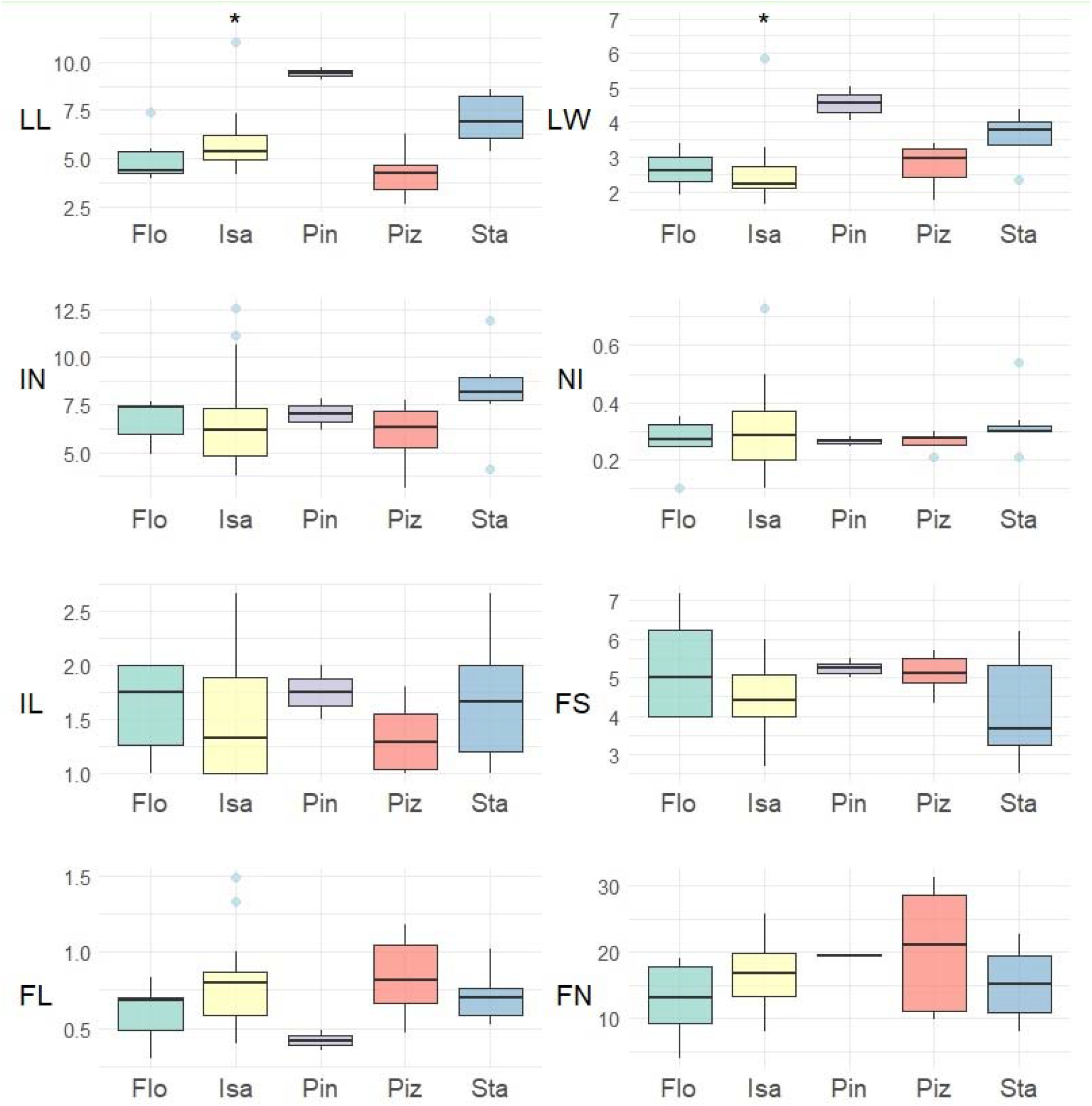
Boxplots showing comparisons of *Phoradendron berteroanum* morphometric traits between islands: Floreana(Flo), Isabela (Isa), Pinta (Pin), Pinzón (Piz) and Santa Cruz (Sta). Asterisks (*) indicate statistically significant differences (p < 0.001).

Morphometric traits are analyzed in each land cover class using an ANOVA test and it was found that two traits are significantly larger in covers dominated by invasive species (LL *p* = 0.00884) and (FL *p* =0.0352). Additionally, the trait IN presents higher values in the land cover ‘Humid grasslands’ (*p* = 0.000394) (Figure 6). It should be noted that some land cover categories are represented by very few specimens, which may impact the reliability of the observed patterns. This limitation must be considered when interpreting the significance of the results, particularly in those categories with minimal sample representation.

**Figure 6.**
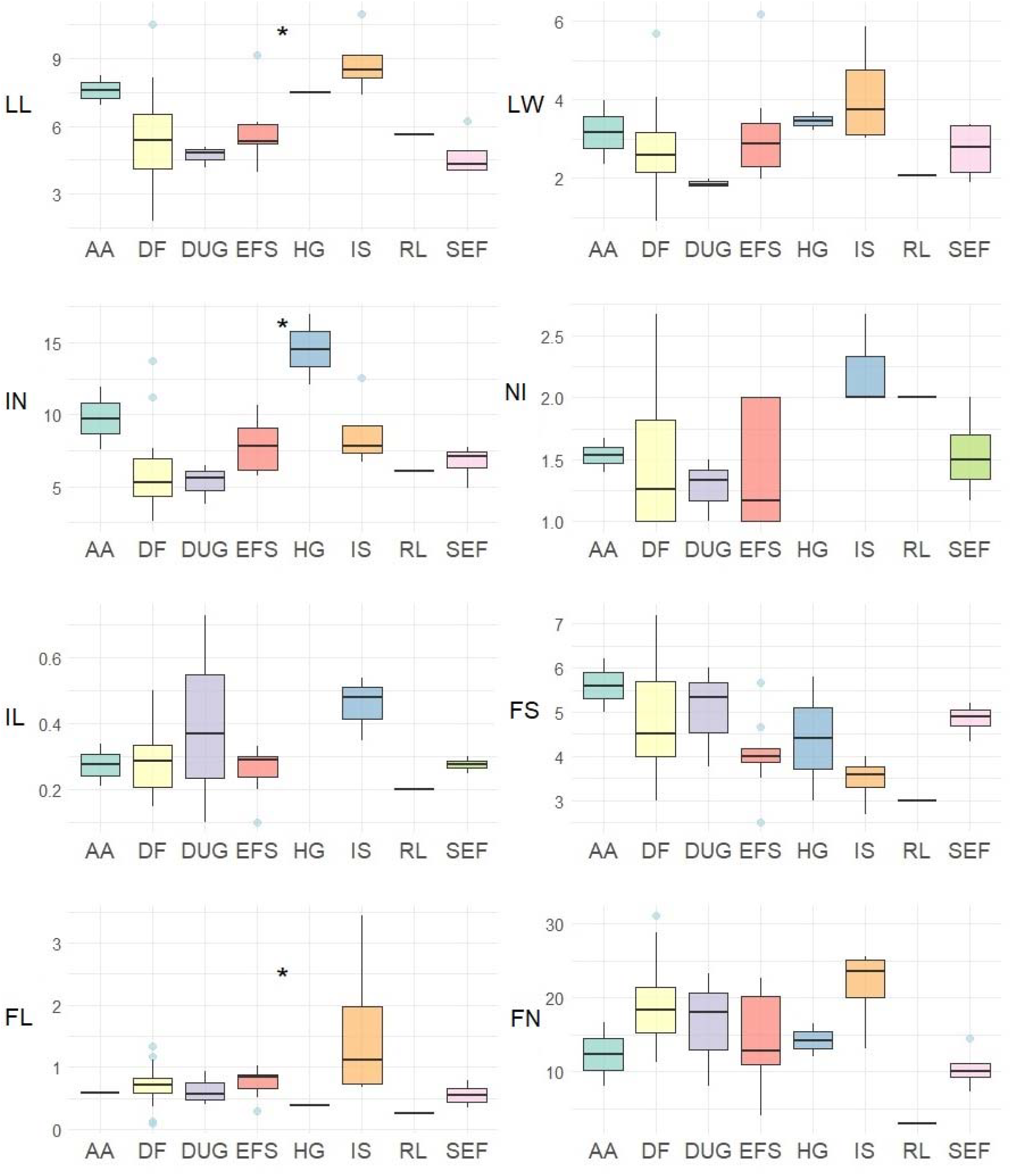
Boxplots showing comparisons of *Phoradendron berteroanum* morphometric traits among the different vegetation covers. Agricultural Area (AA), Deciduous Forest (DF), Deciduous Upland Grassland (DUG), Evergreen Forest and Shrubland (EFS), Humid Grassland (HG), Invasive Species (IS), Recent Lava (LR), Seasonal Evergreen Forest (SEF). Asterisks (*) indicate statistically significant differences (p < 0.001).

### Host specificity

In relation to the hosts, thirty-five parasitism events are recorded on *Phoradendron berteroanum*, with 10 hosts taxa in total. *Zanthoxylum fagara* (L.) Sarg. was the most parasitized species with 14 records (Table 1). The host specificity index (KQ = 0.36) shows a moderate tendency of *P. berteroanum* to parasitize *Z. fagara* in the Galápagos. KQ was also calculated for each elevation group and it was found that for low elevations (KQ = 1) the species exhibits a more generalist tendency compared to medium (KQ = 0.33) and high (KQ = 0.60) elevations. In addition, *Phoradendron berteroanum* produces a higher number of flowers when associated with *Scalesia baurii* B.L.Rob. & Greenm. (FN *p* = 0.00558) (Figure 7).

**Table 1.** Parasitism records and hosts for *P. berteroanum* across three altitudinal ranges: Low (0-499 masl), Medium (500-999 masl) and High (above 1000 masl). Host specificity index was calculated for the species: KQ = 0.36 and for each altitudinal range: KQ_L_ = 1, KQ_M_ = 0.33, and KQ_H_ = 0.60.

| Host species | Altitude range |  |  | Total records |
| --- | --- | --- | --- | --- |
|  | Low | Medium | High |  |
| *Bursera graveolens* | 1 | 0 | 0 | 1 |
| *Croton scouleri* | 1 | 0 | 0 | 1 |
| *ground* | 0 | 0 | 5 | 5 |
| *Lippia rosmarinifolia* | 0 | 0 | 3 | 3 |
| *Trigonopterum laricifolium* | 1 | 0 | 0 | 1 |
| *Psidium* sp. | 1 | 0 | 0 | 1 |
| *Scalesia baurii* subsp. *baurii* | 0 | 3 | 0 | 3 |
| *Scalesia cordata* | 2 | 3 | 0 | 5 |
| *Scalesia pedunculata* | 0 | 1 | 0 | 1 |
| *Zanthoxylum fagara* | 5 | 4 | 5 | 14 |
| Total |  |  |  | 35 |

**Figure 7.**
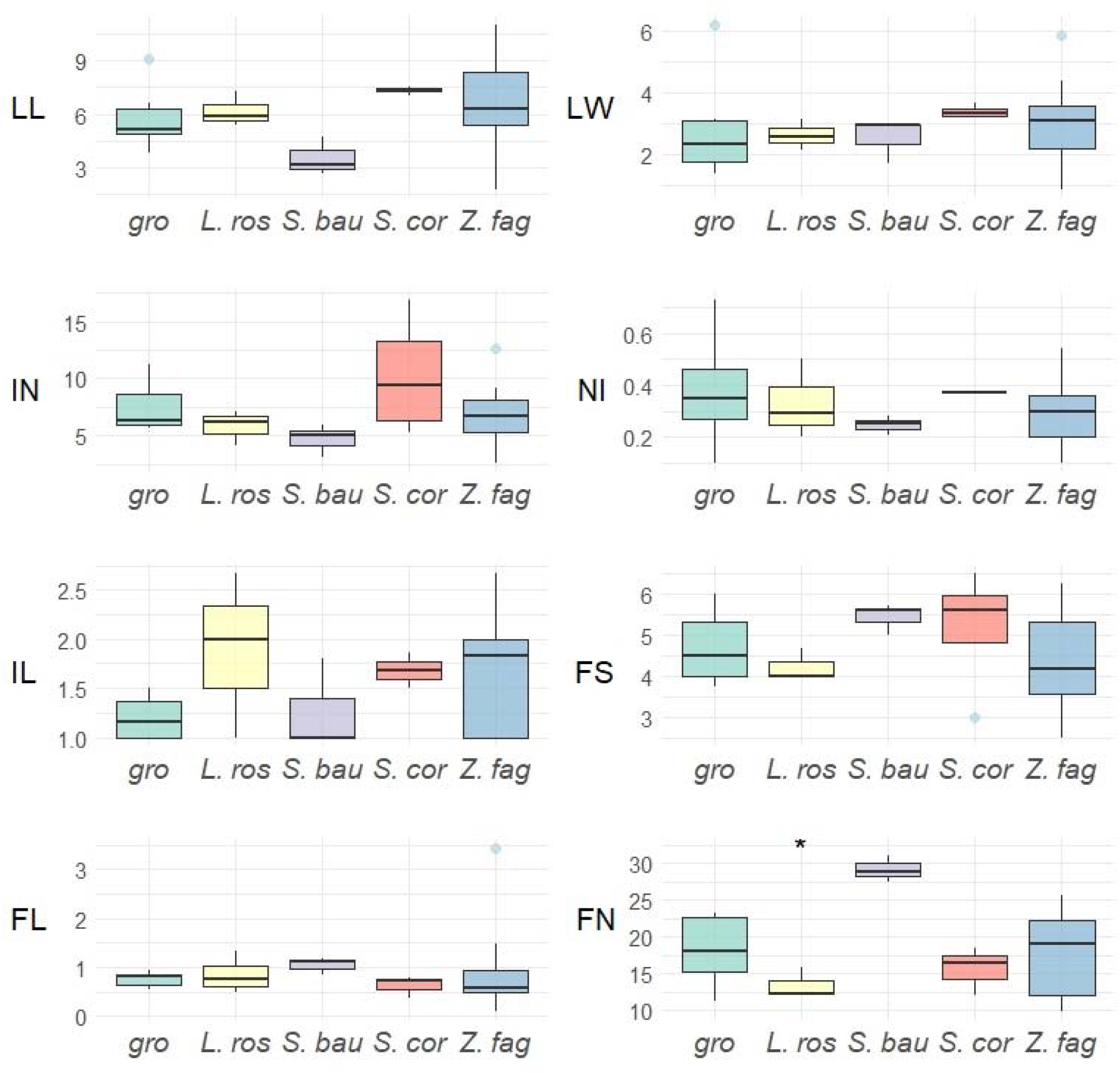
Boxplots showing comparisons of *Phoradendron berteroanum* morphometric traits among recorded hosts: ground (gro*), Lippia rosmarinifolia* (L. ros), *Scalesia baurii* (S. bau), *Scalesia cordata* (S. cor), *Zanthoxylum fagara* (Z. fag). Asterisks (*) indicate statistically significant differences (p < 0.001).

## Discussion

Morphological variation of *Phoradendron berteroanum* could not be fully explained by the altitudinal gradient of the Galápagos Islands. However, significant differences were observed in leaf length (LL), leaf width (LW), and internode length (IN), with the largest values occurring at mid-elevations. Conversely, the number of floral segments (FS) decreased with altitude. Leaf traits (LL and LW) also varied significantly between islands. Notably, LL and flower length (FL) were higher in vegetation covers dominated by invasive species, while IN was greater in humid grasslands. Additionally, flower number (FN) varies significantly by host species; particularly when parasitizing *Scalesia baurii*.

The pronounced altitudinal gradient in the Galápagos Islands shapes vegetation patterns, with arid lowlands and humid highlands differing markedly in precipitation, especially on windward volcanic slopes (Snell & Rea, 1999, Itow, 1992, 2003; Larrea & Di Carlo, 2011). The dry lowlands, which cover 83% of the archipelago, face extreme water scarcity, interrupted mainly by El Niño events (Snell & Rea, 1999). This climatic heterogeneity may influence the physiological responses of plants and drive trait adaptations across elevation zones. For instance, reduced leaf area in *Croton scouleri* Hook.f. at lower elevations mitigates water loss (Castillo et al., 2012), while *Opuntia echios* var. *gigantea* (J.T.Howell) D.M.Porter shows elevation-driven changes in size and reproductive traits (Helsen et al., 2009).

Studies on mistletoes have similarly documented altitudinal variation in fruit morphology, such as larger fruits at lower elevations (Amico et al., 2023). These patterns align with our finding that leaf size increases at mid-elevations in *Phoradendron berteroanum*, suggesting a response to environmental gradients. Comparative data from other mistletoe species support the role of local conditions in shaping morphology, such as geographic differences in the size of *Arceuthobium hondurense* Hawksw. & Wiens (Mathiasen et al., 2012). Such examples suggest that mistletoe morphology is often shaped by environmental variation, although the extent and nature of this plasticity may differ across taxa. These patterns reinforce the view that both abiotic conditions and potential local adaptation play a role in mistletoe trait variability.

Parasitic plants, like mistletoes, rely on hosts for water and nutrients (Kuijt, 1969; Phoenix & Press, 2005; Watson et al., 2022). Their high transpiration rates and limited stomatal control allow passive water uptake under normal conditions, but increase vulnerability to drought stress for both parasite and host (Griebel et al., 2022; Henríquez□Velásquez et al., 2012; Watson et al., 2022). In dry habitats such as the Galápagos lowlands, reduced leaf size likely reflects an adaptive strategy to minimize water loss, consistent with reports in other mistletoe species under water stress (Phoenix & Press, 2005). The greater number of floral segments observed at lower elevations may represent a shift in resource allocation toward reproduction under stressful conditions, as seen in other taxa (Nabity et al., 2021; Sala et al., 2001).

*Phoradendron berteroanum* is broadly distributed across the Galápagos occupying diverse habitats from arid lowlands to high-altitude grasslands, including old lava fields, *Scalesia* forests, and littoral zones. Its prevalence across disturbed and undisturbed vegetation types reflects ecological resilience. At mid-elevations—where temperature and moisture conditions may be more favorable—the species displays increased leaf and internode size, potentially enhancing photosynthetic efficiency and internal transport. (Leigh et al., 2017; Okajima et al., 2012; Parkhurst and Loucks, 1972)

Also, these traits may indicate strategic resource investment in vegetative growth, consistent with the phenotypic plasticity reported in other mistletoe species (Watson et al., 2022). Additionally, significant differences in LL and LW between islands further support the influence of local environmental factors, particularly altitude and water availability. For instance, Isabela’s broad elevational and climatic range appears to favor larger leaves, while Pinta’s lower and drier conditions favor smaller leaves as an adaptation to increased water stress (Rivas-Torres et al., 2018; Trueman, 2010). However, it is important to note that the sample size for Pinta remains disproportionately small compared to other islands, with only a handful of specimens available. This limitation must be borne in mind when interpreting any significant differences involving Pinta, as the scant representation undermines the robustness of island□level comparisons.

It is recommended incorporating additional environmental factors beyond elevation. First, the moisture regime differs dramatically between windward and leeward slopes of the Galápagos volcanoes—windward slopes receive far more rainfall from the trade winds than their leeward counterparts. Thus, mean precipitation may be a more informative predictor of leaf size than altitude alone (Jefferson et al., 2014; Mouginis□Mark et al., 1996). Second, cloud cover patterns also vary with topography: the highland humid zone is characterized by persistent clouds, whereas the adjacent pampa belt—despite occurring at similar or even higher elevations— experiences much greater insolation (Pryet et al., 2012). Accounting for these cloud dynamics could reveal how reduced light availability drives the evolution of larger leaves to optimize photosynthetic capacity. Integrating these factors into forthcoming analyses will be crucial to untangling the true environmental drivers of leaf morphology in *Phoradendron berteroanum*.

Taken together, these findings highlight the influence of environmental heterogeneity on the phenotypic plasticity of *Phoradendron berteroanum* across the Galápagos Islands. While lowland populations exhibit traits that minimize water loss and enhance reproductive output, mid-elevation populations emphasize vegetative growth and resource acquisition. Comparisons between Pinta and Isabela reinforce the importance of localized environmental drivers in shaping growth and reproductive strategies. Nonetheless, the small sample size from Pinta warrants caution in making broader generalizations about the morphological variation between islands.

Regarding host specificity, *Phoradendron berteroanum* exhibits a moderate preference for *Zanthoxylum fagara*, yet overall parasitizes a wide range of hosts, particularly at lower elevations (KQ = 1). This pattern is consistent with evidence that *Phoradendron* parasitizes families from phylogenetically distant orders, including Sapindales, Rosales, Gentianales, Lamiales, Ericales, Myrtales, Malvales, Caryophyllales, and others, confirming its broad taxonomic and phylogenetic host range (Tinoco□Domínguez et al., 2024). Moreover, although some mistletoe species—such as *Phoradendron californicum*—primarily parasitize host species from a single family (e.g., Fabaceae), not all *Phoradendron* species display such host specificity at the family or functional group level (e.g., nitrogen-fixers). Assuming uniformity at the genus level may therefore lead to overgeneralization, distort interaction networks, inflate perceived host ranges, or underestimate parasite specialization (Tinoco□Domínguez et al., 2024).

Higher host specificity at mid- and high-elevations may reflect differences in host availability or physiological compatibility. Moreover, when growing on *Scalesia baurii, Phoradendron berteroanum* produces significantly more flowers, suggesting that host species influences reproductive output—perhaps through differential nutrient uptake or elevation-linked host traits (Westwood et al., 2010). Herbarium specimens provide a valuable historical record for assessing how *P. berteroanum* responds to environmental heterogeneity across islands and elevations. By combining these archival data with ecological observations, a solid picture of the phenotypic plasticity and adaptive strategies of the species is obtained. Nonetheless, several limitations must be acknowledged. The sample size is small (68 specimens across eight islands), and its distribution is uneven. When divided by island, elevation range, or land cover class, some categories include only one or very few specimens, which compromises statistical comparisons. Additionally, sampling bias is likely, as many specimens were collected opportunistically—often from plants that were more accessible—potentially underrepresenting the full structural variability of the parasite. Finally, working with dry herbarium material can obscure key traits, especially those related to leaf texture or three-dimensional structure. However, this bias is partially mitigated by the fact that all specimens underwent similar preservation protocols, making comparisons among them still informative.

Overall, morphological variation in *Phoradendron berteroanum* in the Galápagos cannot be attributed solely to altitudinal gradients but rather to a complex interplay of water availability, island vegetation type, and host compatibility. Mid-elevation populations develop larger leaves and longer internodes—traits likely enhancing photosynthetic capacity and internal transport— whereas lowland populations bear smaller leaves and more floral segments, a reproductive strategy suited to water-limited conditions. Contrasting patterns between islands such as Isabela and Pinta underscore the influence of local microclimates and elevation ranges, while host-specific differences in fecundity (e.g., on *Scalesia baurii*) highlight the critical role of parasite– host interactions in shaping the ecology of this mistletoe. To deepen our understanding of these patterns, future research should move beyond elevation as a sole explanatory variable. While altitude serves as a general proxy for climate, it does not account for key environmental gradients across the archipelago. For instance, the windward slopes of Galápagos volcanoes receive significantly more precipitation due to the prevailing trade winds, while leeward slopes remain markedly drier (Jefferson et al., 2014; Mouginis□Mark et al., 1996). Additionally, cloud cover patterns vary with topography: persistent cloudiness in the highland humid zone reduces solar radiation, whereas the adjacent pampa belt—often at comparable or even higher elevations— experiences much greater insolation (Pryet et al., 2012). Including spatial data on rainfall and cloud cover in future analyses could help determine whether observed morphological traits, such as larger leaves, represent adaptations to light-limited rather than elevation-related environments. This integrative approach will be essential for identifying the true environmental drivers of phenotypic variation in *P. berteroanum*.

This study emphasizes the value of herbarium specimens for morphological research, extending their utility beyond traditional fields such as taxonomy, systematics, and evolutionary studies (Davis, 2023). Herbaria are increasingly being recognized as valuable resources for broader ecological investigations, including the study of phenology, phenotypic plasticity, and plant responses to climate regimes or climate change (Meineke et al., 2018; Swain and Chakraborty, 2024). This is especially the case when such investigations are combined with emerging and increasingly accessible genomic techniques; however, it is important to acknowledge the inherent limitations of herbarium-based trait studies. The opportunistic nature of specimen collection can introduce biases; for instance, botanists often prioritize larger, more conspicuous, or more accessible individuals, and typically only collect parts of the plant, particularly in woody species. Consequently, trait data may not accurately represent the full morphology of the individual or the variability within populations (Heberling, 2022; Willis et al., 2017). Furthermore, the presence of duplicate specimens and uneven sampling across taxa, regions, and time periods has the potential to compromise the accuracy and representativeness of trait-based analyses derived from herbarium collections (Willis et al., 2017).

## Conclusions

Although this study did not detect consistent significant differences across elevation zones, islands, host species, or vegetation types, certain morphological traits—particularly those related to leaf size and inflorescence structure—exhibited variation in response to local environmental conditions. These patterns suggest high phenotypic plasticity in *Phoradendron berteroanum*, likely driven by the environmental heterogeneity characteristic of the Galápagos Archipelago.

Specifically, the increase in leaf size at mid-elevations and in humid environments, along with a reduction in reproductive structure size in these same areas, may reflect an ecological strategy influenced by water availability. In wetter habitats, where water deficit is less severe, the species’ limited stomatal control may pose less of a physiological constraint. In contrast, in arid lowland areas, reduced leaf size and increased reproductive investment—evidenced by a greater number of flowers and smaller leaves—suggest a shift toward maximizing reproductive output under hydric stress.

The integration of herbarium-based morphological data with ecological context provides valuable insights into the adaptive strategies of *Phoradendron berteroanum*. This approach contributes to a broader understanding of how parasitic plants respond to environmental gradients and may inform predictions about their responses to future climate change in insular ecosystems.

## Acknowledgements

We thank Nicolás Moity for critically reading this paper. The data for this research was obtained when AD participated as volunteer at the Charles Darwin Foundation. Thanks to the Directorate of the Galápagos National Park (DPNG) and the Ministry of Environment (MAATE), for providing us with the permits for the annual operation of the Galápagos Natural History Collections (N° MAATE-MCMEVS-2024-075) under which this project was conducted. This publication is contribution number 2737 of the Charles Darwin Foundation for the Galápagos Islands

## Credit Statement (for author contributions)

Juliana Puentes-Marín (Conceptualization, Data curation, Formal analysis, Visualization, Writing – original draft), Anakah Denison (Data acquisition, Writing – review & editing), Justine Villalba-Alemán (Data acquisition, Writing – review & editing), Johny Mazón-Redín (Visualization), and C. Miguel Pinto (Conceptualization, Writing – review & editing).

## Supporting information

The following supporting information can be downloaded at:

Puentes-Marín, J., Denison, A., Villalba-Alemán, J., Mazón-Redín, J., & Pinto, C. M. (2025). Morphological and ecological data for Phoradendron berteroanum specimens deposited in CDS collection. [Data set]. rdm.https://researchdata.fcdarwin.org.ec/records/202cw-xhy96 **Table S1**. Morphological and ecological data for *Phoradendron berteroanum* specimens deposited in CDS collection.

## Data Availability Statement

The data that support the findings of this study are available at: https://researchdata.fcdarwin.org.ec/records/202cw-xhy96

